# Mutant p53 Misfolding and Aggregation Precedes Transformation into High-Grade Serous Ovarian Carcinoma

**DOI:** 10.1101/2024.09.17.612958

**Authors:** Sara Sartini, Lexi Omholt, Neda A. Moatamed, Alice Soragni

## Abstract

High Grade Serous Ovarian Cancer (HG-SOC), the most prevalent and aggressive gynecological malignancy, is marked by ubiquitous loss of functional p53, largely due to point mutations that arise very early in carcinogenesis. These mutations often lead to p53 protein misfolding and subsequent aggregation, yet the alterations in intracellular p53 dynamics throughout ovarian cancer progression remain poorly understood. HG-SOC originates from the fallopian tube epithelium, with a well-documented stepwise progression beginning with early pre-malignant p53 signatures. These signatures represent largely normal cells that express and accumulate mutant p53, which then transform into benign serous tubal intraepithelial lesions (STIL), progress into late pre-malignant serous tubal intraepithelial carcinoma (STIC), and ultimately lead to HGSOC. Here, we show that the transition from folded, soluble to aggregated mutant p53 occurs during the malignant transformation of benign precursor lesions into HGSOC. We analyzed fallopian tube tissue collected from ten salpingo-oophorectomy cases and determined the proportion of cells carrying soluble versus mis-folded/mutant p53 through conformation-sensitive staining and quantification. Misfolded p53 protein, prone to aggregation, is present in STICs and HG-SOCs, but notably absent from preneoplastic lesions and surrounding healthy tissue. Overall, our results indicate that aggregation of mutant p53 is a structural defect that distinguishes preneoplastic early lesions from late premalignant and malignant ones, offering a potential treatment window for targeting p53 aggregation and halting ovarian cancer progression.

## Introduction

p53 is the critical tumor suppressor^1,2^, and responds to stress signals due to DNA damage, hypoxia or oncogenic signaling by arresting the cell cycle and inducing either DNA repair or apoptosis^3^. By tightly regulating the cell cycle, p53 plays an essential role in preventing cancer progression, preventing cells from growing and dividing too fast in an uncontrolled manner. In normal conditions, p53 levels are strictly regulated by the interaction with MDM2, a E3 ubiquitin ligase, which leads to rapid proteasomal degradation of p53^4^. Half of all human cancers lose p53 function by missense mutations, which result in the loss of DNA binding capacity or in the destabilization of the protein toward a partially unfolded conformation^5–7^. The mutations that destabilize p53 structure cause the exposure of the hydrophobic adhesive core of the protein, including the p53_252-258_ segment which rapidly interacts with other p53 molecules causing aggregation^8–12^. Mutant p53 aggregates exert a dominant-negative effect by sequestering and inactivating wild type p53, thereby blocking its tumor suppressive transcriptional activity, apoptotic functions and interaction with MDM2, ultimately leading to protein accumulation^9–11^. Beyond diminishing its tumor suppressor role, recent studies have shown that p53 aggregation can actively enhance the transformation of cells into cancer^12,13^. These aggregates also exhibit gain-of-function properties, promoting alterations in cell cycle regulation and proliferation pathways, chromatin reorganization, and resistance to senescence^12,13^. Additionally, p53 aggregation has been associated with the upregulation of cell metabolism, chaperone machinery, and the unfolded protein response^12,13^. Tumors harboring aggregated mutant p53 tend to respond poorly to chemotherapy and are associated with worse clinical outcomes^14–16^.

High-grade serous ovarian carcinomas (HGSOCs), the most common and deadly type of ovarian cancer, are a prime example of tumors biting p53 aggregation^9,17^. HGSOC is characterized by the widespread loss (96%) of functional p53, primarily due to missense mutations that occur early in carcinogenesis. These mutations often result in p53 misfolding and aggregation in advanced stages of HGSOC, as we and others have previously demonstrated^9,18–21^. HGSOCs are aggressive tumors that are often diagnosed at advanced stages, when the disease has already spread to distant sites^19,22^, resulting in a five-year survival rate of 30.8%^22^. However, when detected early and still localized to the primary site (stage I), the five-year survival rate is significantly higher (93.1%)^22^. One of the most significant challenges to improving patient survival is the lack of effective screening tests for early detection and monitoring of ovarian cancer progression. Although progress has been made in developing new treatments for HGSOCs in recent years, including the approval of the folate receptor-targeting antibody-drug conjugate mirvetuximab soravtansine-gynx^23^, the backbone of therapy remains Carboplatin/Taxol combination in the first line setting. Despite high initial response rates, the majority of patients eventually experience disease recurrence, which remains hard-to-treat^24,25^.

HGSOC is now understood to originate from the fallopian tube epithelium rather than the ovary^26^. It develops from early precursor lesions known as serous tubal intraepithelial carcinoma (STIC)^26–31^. These noninvasive lesions are marked by features such as hyperchromasia, nuclear atypia, loss of cell polarity, increased proliferative activity, and a high frequency of p53 mutations^32,33^. STICs are frequently identified in the fallopian tubes of BRCA mutation carriers undergoing preventive salpingo-oophorectomy and are also a concurrent finding in approximately 60% of sporadic HGSOC cases^27,29,34^. In addition to STICs, p53 mutations have been observed in pre-neoplastic cell populations within the benign tubal mucosa, referred to as p53 signatures^35,36^. These early lesions are composed of clusters of 10-30 benign-appearing epithelial cells with low proliferative activity and upregulated mutated p53 expression^36,37^. Another lesion type, termed serous tubal intraepithelial lesion (STIL), represents an intermediate stage between early p53 signatures and more advanced STICs^38^. STILs exhibit over-expression of mutated p53 in a larger number of cells compared to p53 signatures, along with increased proliferative activity, though not to the extent seen in STICs^38,39^. Both p53 signatures and STIC lesions are found in the distal end of the fallopian tube^26,37^, and as such, p53 signatures appear to precede STIC formation and arise early in carcinogenesis^26^. Most importantly, STIC and HG-SOC from the same patient typically share the same p53 mutations, supporting a possible clonal relationship in many cases^26–28^.

Mutations in p53 are recognized as major drivers of HGSOCs, yet these mutations occur early, before malignant transformation, leaving the mechanisms underlying tumor development unclear. p53 mutations can lead to protein aggregation and functional inactivation. Our previous studies have shown that p53 aggregates are present in HGSOCs, and that the inhibitor peptide ReACp53 can target and prevent this aggregation, leading to reactivation of p53 and cell death^9^. However, the timing of p53 aggregation and its potential role in driving the progression from early precursor lesions to malignant tumors remain uncertain. In this study, we explored the presence of mutant p53 aggregation across different stages of disease progression, including p53 signatures, STILs, STICs, and HGSOCs in clinical samples. Our findings suggest that p53 aggregation is a distinct structural defect that differentiates preneoplastic lesions from premalignant and malignant ones. This distinction highlights a potential window for early intervention and prevention of ovarian cancer progression.

## Results

### Patient characteristics and p53 mutational status

To detect p53 aggregation at various stages of HGSOC progression, we obtained tissue samples from n=10 patients who underwent either salpingo-oophorectomy as part of preventive care (for BRCA carriers) or debulking surgery for HG-SOC (**Table 1**). The mean age of the patients was 60 years old, ranging from 45 to 74. Each sample was carefully examined for the presence of four types of lesions. Patients CA4 to CA9 exhibited a complete range of lesions, from p53 signatures to HGSOC. Patients CA1 and CA10 had p53 signatures, STILs, and STICs, while patient CA2 showed only advanced disease, specifically STICs and HGSOCs. Patient CA3 did not have any detectable abnormalities in the sections received and was excluded from further analysis. The specific findings for each patient are detailed in **Supplementary Table 1**.

**Table 1.** List of patients and tumor sample characteristics.

| Patient ID | Age* | Diagnosis** | Histological grade | Specimen |
| --- | --- | --- | --- | --- |
| CA 1 | 60 | High grade serous papillary carcinoma | N/A | Fallopian tube with tubal intraepithelial carcinoma |
| CA 2 | 50 | High grade serous adenocarcinoma with lymphovascular invasion | NA | Fallopian tube with serous tubal intraepithelial carcinoma |
| CA 3 | 50 | High grade serous papillary carcinoma in ovarian cyst | G3 | Fallopian tube with tubal intraepithelial carcinoma and paratubal cyts |
| CA 4 | 60 | High grade papillary serous adenocarcinoma | G2 | Fallopian tube with adenocarcinoma and tubal intraepithelial carcinoma |
| CA 5 | 50 | High grade serous carcinoma | G3 | Adnexa, fimbriae and tumor with tubal intraepithelial carcinoma |
| CA 6 | 40 | High grade serous carcinoma | G3 | Fallopian tube and fimbria with invasive carcinoma and serous tubal intraepithelial carcinoma |
| CA 7 | 70 | High grade serous carcinoma | G3 | Fallopian tube with tubal intraepithelial carcinoma |
| CA 8 | 70 | Serous carcinoma with tubal intraepithelial carcinoma | NA | Fallopian tube with tubal intraepithelial carcinoma |
| CA 9 | N/A | High grade papillary serous adenocarcinoma | NA | Fallopian tube with serous tubal intraepithelial carcinoma |
| CA 10 | N/A | High grade serous carcinoma with extensive necrosis | NA | Adnexa, fallopian tube with serous tubal intraepithelial carcinoma |

\*Decade of age at time of tissue procurement. \*\*Diagnosis at time of tissue procurement. N/A information not available

To confirm that p53 mutations occur early in HGSOC development and that p53 aggregation may be a transitional event in the progression of ovarian cancer, we selected representative lesions from each patient for sequencing. Due to the small size of pre-neoplastic lesions, we pooled several areas to obtain sufficient DNA for sequencing, while larger, more advanced lesions were sequenced individually. We successfully obtained representative sequencing results for three patients: CA2 for late-stage lesions, CA10 for early-stage lesions, and CA9 for lesions spanning the full range of benign to malignant (**Figure 1**). In all three samples, p53 harbored somatic missense mutations in the DNA-binding domain, a region frequently mutated and associated with protein misfolding^6,9,40,41^. Notably, p53 mutations in patient CA9 were identical across early and late lesions, including p53 signatures and HGSOC, indicating that the mutations themselves may not drive tumor progression. This finding supports further investigation into the conformational status of p53 as a potential critical factor in disease progression.

**Figure 1.**
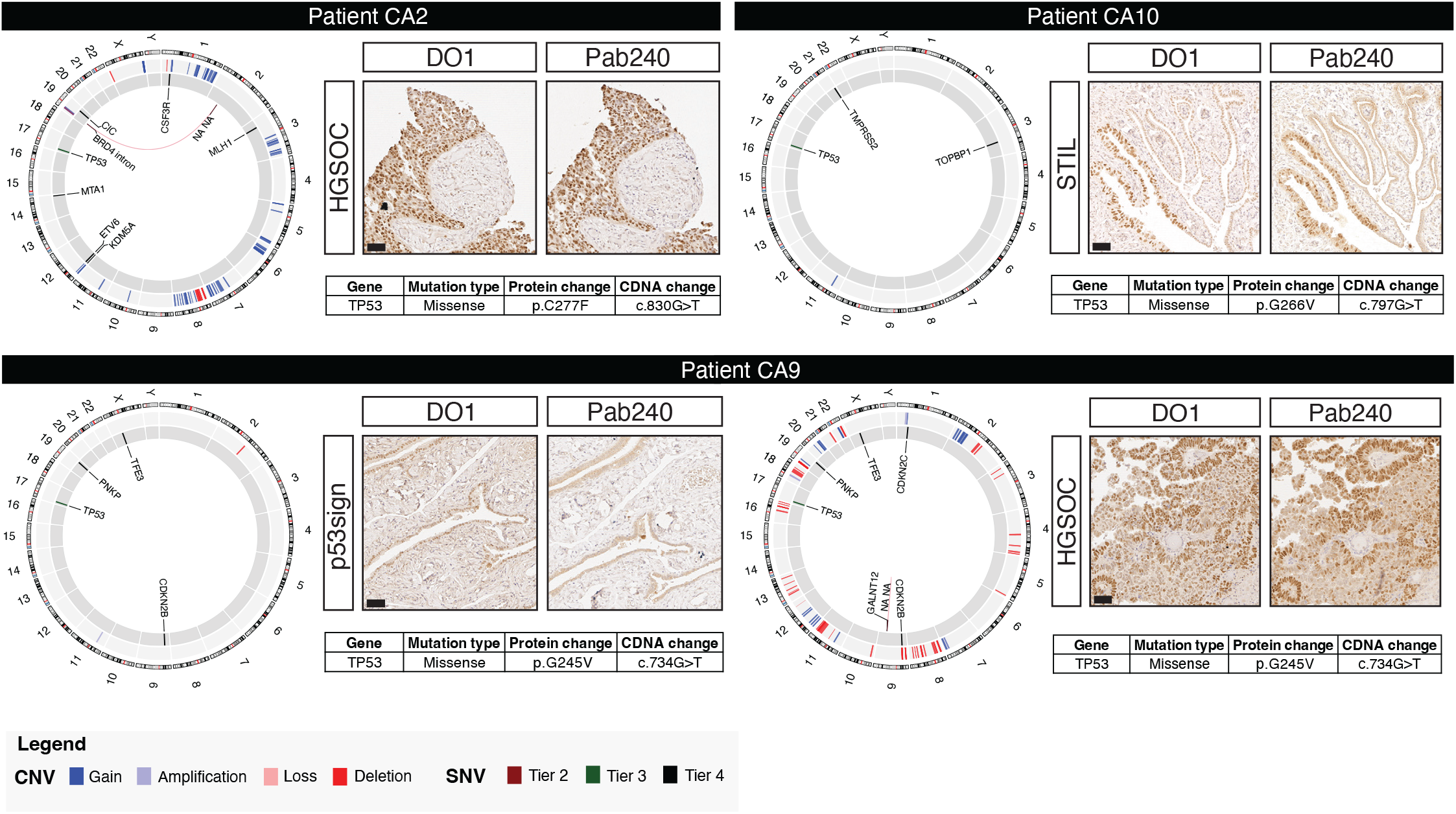
Mutation status of TP53 in representative patients. Sequencing results confirmed mutations in TP53 from the early stages of the tumor. Circos plots show all mutations and structural rearrangement found in each sample. Representative area used for sequencing is shown on the right of the plot and table reporting the mutation found in TP53 gene and the consequent aminoacidic change. Here is shown three representative patients with late-stage lesions (CA2), early-stage lesions (CA10) and both early and late stages lesions (CA9). Scale bar 50 µm. p53sign, p53signature; HGSOC, high-grade serous ovarian cancer.

### p53 protein conformation in pre-malignant and malignant samples

For each patient, we stained paraffin-embedded tissue sections with hematoxylin and eosin (H&E) and performed immunohistochemistry. We used the pan-p53 antibody DO-1, which detects all forms of p53 regardless of folding status, and the PAb240 antibody, which specifically identifies misfolded p53, a precursor to p53 aggregation^9,42^. PAb240 binds to an epitope (aa 213-217) which is buried when the protein is properly folded and exposed upon mutation and unfolding^9,42^. Additionally, we stained the sections for Ki-67 to assess the proliferative activity of the tissue.

After identifying areas in the histological sections that contained p53 signatures, STILs, STICs and HGSOC, we used DO-1 immunostaining to assess the upregulation of p53 in each lesion. A pathologist verified appropriateness of the identified areas before further analysis. We compared the expression level of p53 detected by DO-1 with those detected by Pab240, to understand which lesions were associated with p53 misfolding as a surrogate marker of aggregation. All the lesions exhibited high levels of p53 accumulation, including p53 signatures, STILs, STICs and HGSOC areas. However, misfolded p53 was only present in STICs and HGSOCs, and notably absent from p53 signatures and STILs as well as surrounding normal tissue (**Figure 2 and Supplementary Figures 1-7**). **Figure 2** highlights representative areas from patients CA4 and CA8, for which we could identify the full range of benign to malignant lesions within the same section. Representative areas for the other patients are shown in **Supplementary Figures 1-7**. Ki-67 staining, which indicates proliferative activity, was either absent or low in p53 signatures and STILs, whereas STICs and HGSOCs exhibited high proliferative activity, consistent with more advanced stages of cancer and previous literature reports^36,38,43^.

**Figure 2.**
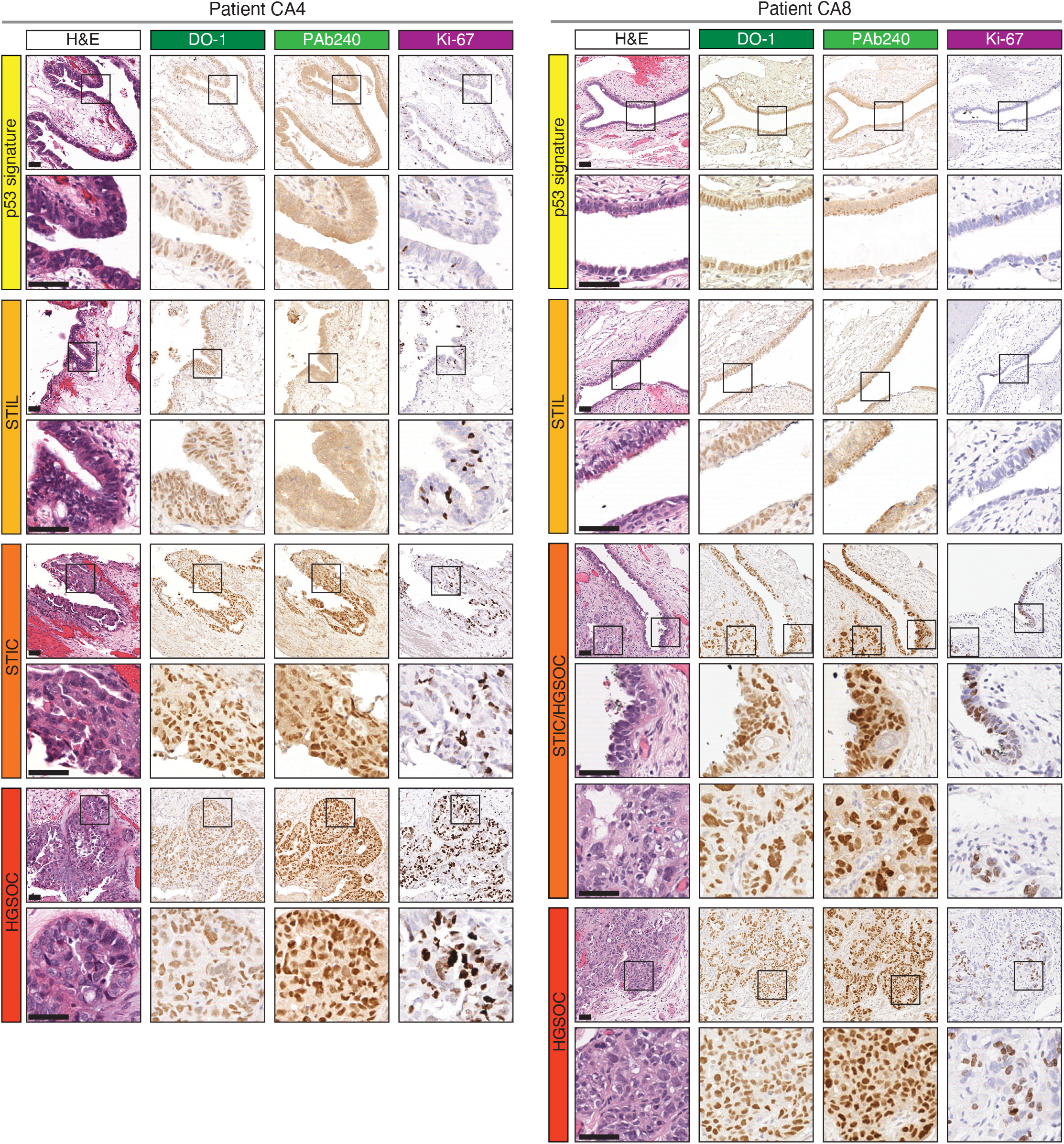
p53 upregulation and aggregation in all four lesion types. Patients CA4 and CA8 exhibit the entire range of lesions: p53 signature, STIL, STIC and HGSOC. Staining with H&E, DO-1, PAb240 and Ki-67 are reported for each representative lesion. Black boxed area is magnified on the bottom. Scale bars, 50 µm, magnification 20 µm.

We also performed staining for the transcrption factor paired box 8 (PAX8), which is known to be highly upregulated in HGSOCs, suggesting its critical role in this tumor type^44^. In our analysis, PAX8 was strongly expressed in late-stage lesions, while only a few positive nuclei were detected in early-stage areas, as shown for representative patients in **Supplementary Figures 8-11**.

### Quantification of p53 aggregation in different stages of HGSOC

To better characterize the extent of p53 misfolding/aggregation observed in our histological examination, we employed an image-based quantification method on DO-1 and PAb240 staining to quantify both total and aggregated p53 protein expression. We applied the same method to quantify Ki-67 expression to assess the proliferation index at different stages of tumor progression. For each patient, the analyzed areas were processed using Fiji’s Macro Tool^45^ to calculate a positive index for each staining, indicative of the number of positively stained nuclei relative to the total number of nuclei in a specific area.

Quantification of p53 levels in patients CA4 and CA8 (DO-1, **Figure 3**) revealed a comparable expression level across all lesion types, confirming that p53 upregulation is a very early phenomenon that precedes ovarian cancer progression. However, misfolded/aggregated p53 as detected by PAb240 was only found in STICs and HGSOC. We calculated the cumulative positive index for DO-1, PAb240, and Ki-67 for all lesions collected from these patients (**Figures 3A and 3C**). The ratio of PAb240 to DO-1 in each area was close to 1 in STICs and HGSOCs (**Figures 3B and 3D, Supplementary Table 2**), indicating that nearly all p53 present at these stages is misfolded/aggregated.

**Figure 3.**
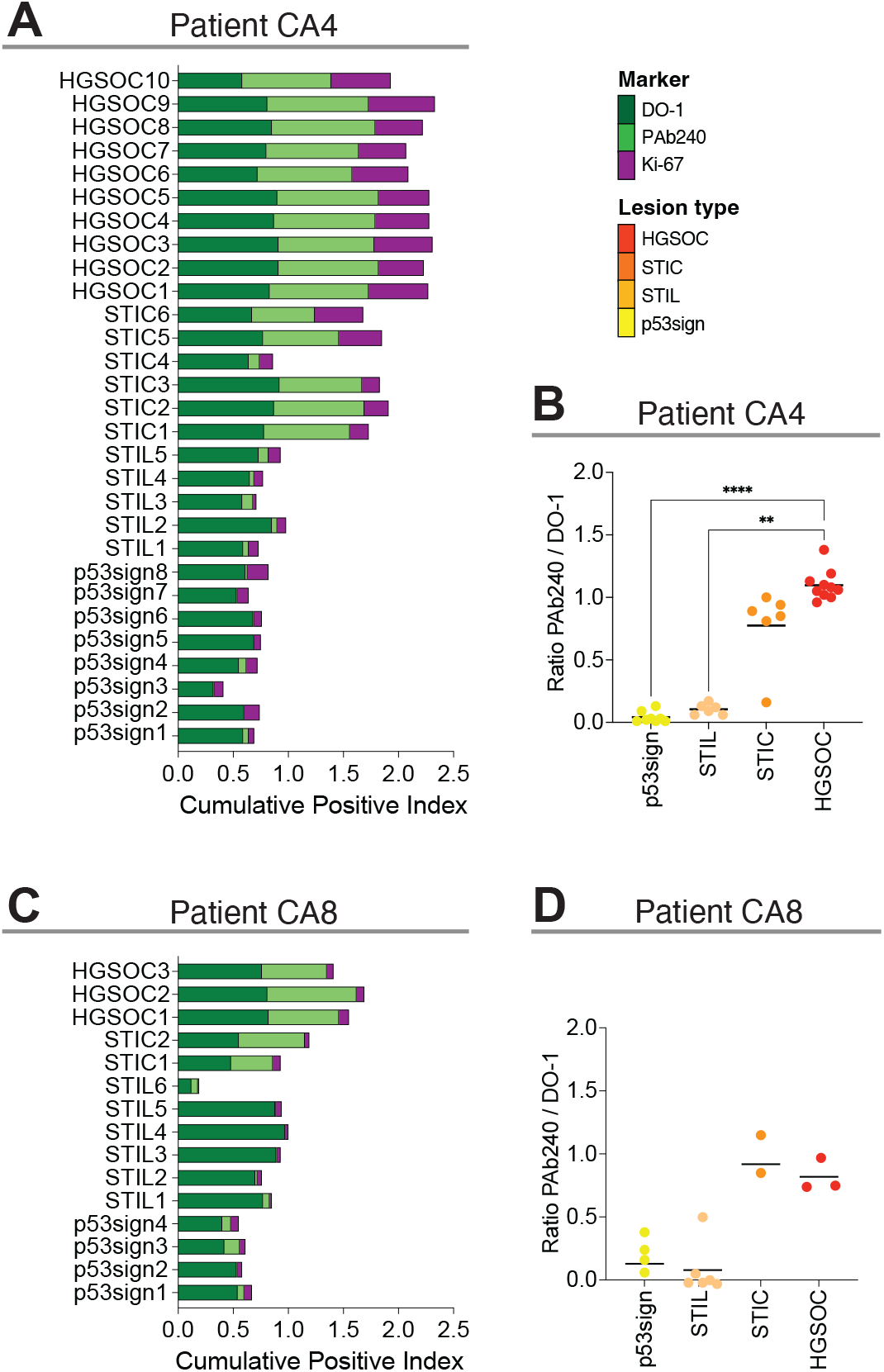
Quantification of total and aggregated p53 in repre-sentative patients. (A and C) Positive index calculated for DO-1 (dark green), PAb240 (light green) and Ki-67 (purple) is reported for each lesion collected for patients CA4 and CA8. Each bar represents a lesion. The graphs report the cumulative positive index and show that the aggregation of p53 is present only in late stages of the tumor progression. (B and D) Ratio of Pab240 and DO-1 calculated for each lesion (p53 signature yellow, STILs light orange, STIC dark orange, HGSOC red). Each dot represents the ratio of a single lesion. For Patient CA4 (B): p53sign n=8, STIL n=5, STIC n=6, HGSOC n=10. For Patient CA8 (D): p53sign n=4, STIL n=6, STIC n=2, HGSOC n=3. Statistical significance tested by performing a Kruskal-Wallis test with Dunn’s correction for multiple comparisons. **p < 0.05, ****p < 0.0001; ns are not specified in the graphs. HGSOC, high-grade serous ovarian cancer; STIC, serous tubal intraepithelial carcinoma; STIL, serous tubal intraepithelial lesion; p53sign, p53 signature.

In contrast, this ratio was substantially lower in p53 signatures and STILs, confirming the near absence of p53 aggregates despite high levels of mutated p53. Results observed for patients CA4 and CA8 are consistent with all other patient studies (**Supplementary Figure 12 and Supplementary Table 2**). Combining the findings from all patients analyzed in this study consistently demonstrates a clear trend of increasing p53 aggregation as ovarian cancer progresses. In the early lesions, such as p53 signatures and STILs collected from 9 patients, p53 is largely well-folded while in late lesions STICs and HGSOCs, nearly all detected p53 was misfolded/aggregated (**Figure 4, Supplementary Figure 13 and Supplementary Table 2**). Additionally, both STICs and HGSOCs showed significantly higher Ki-67 indices, indicating increased cell proliferation, with values ranging from 5% to 49.4%, compared to the lower Ki-67 indices observed in p53 signatures and STILs, which ranged from 0.9% to 9.9% (**Supplementary Figure 14**). This correlation suggests that p53 misfolding/aggregation and increased proliferative activity are linked to ovarian cancer progression.

**Figure 4.**
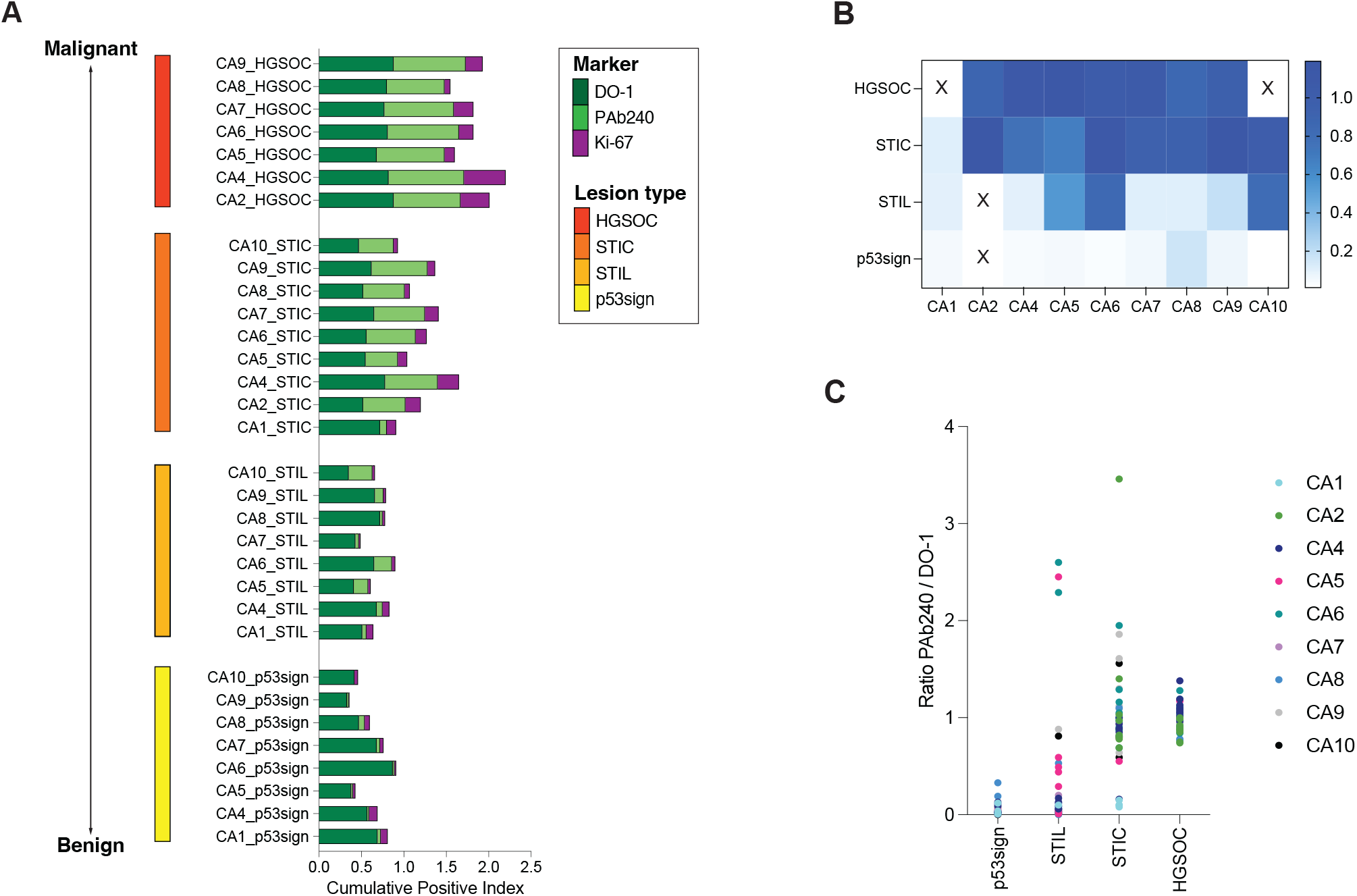
p53 quantification overview in all patients. (A) Cumulative positive index calculated for DO-1 (light green), Pab240 (dark green) and Ki-67 (purple) for each patient, organized by lesion progression from benign to malignant. Each bar represents the quantification of all lesions of a given type found in a patient. In this summary, it is evident how p53 aggregation increases with disease progression. (B) Heatmap of the calculated Pab240/DO1 ratio for all four types of lesions found in each patient. The blue scale shows how the ratio approaches 1 with disease progression, highlighting that all p53 present in that lesion is aggregated. “X” indicates the absence of a lesion type in that specific patient. (C) Ratio of Pab240 and DO-1 calculated for each lesion in each patient. Each dot represents the ratio of a single lesion collected in one patient. CA1: p53sign n= 4, STIL = 1, STIC n= 4; CA2: STIC n= 9, HGOSC n=10; CA4: p53sign n=8, STIL n=5, STIC n=6, HGSOC n=10; CA5 : p53sign n=2, STIL n= 8, STIC n= 2, HGSOC n= 2; CA6: p53sign n= 1, STIL n= 6, STIC n= 10, HG-SOC n= 10; CA7: p53sign n= 3, STIL n= 2, STIC n= 1, HGSOC n=1; CA8: p53sign n=4, STIL n=6, STIC n=2, HGSOC n=3; CA9: p53sign n= 8, STIL n= 5, STIC n= 9, HGOSC n= 10; CA10: p53sign n= 4, STIL n= 1, STIC n=3. HGSOC, high-grade serous ovarian cancer; STIC, serous tubal intraepithelial carcinoma; STIL, serous tubal intraepithelial lesion; p53sign, p53 signature.

The analysis revealed a distinct pattern in ovarian cancer progression: early lesions, such as p53 signatures and STILs, exhibited generally low proliferative activity, as indicated by Ki-67 staining yet displayed high levels of p53, although the protein remained largely well-folded (**Figure 4A and Supplementary Figure 14**). In contrast, late-stage lesions, including STICs and HGSOCs, retained comparably high p53 levels but were characterized by a significant increase in both p53 misfolding/aggregation and cell proliferation (**Figure 4A and Supplementary Figure 14**). This shift from well-folded p53 in early lesions to misfolded and aggregated p53 in advanced stages highlights a critical transition point that may have critical relevance to the progression of ovarian cancer.

## Discussion

In HGSOC, mutations in p53 are ubiquitous and occur very early in disease development. Some of these mutations lead to the misfolding and aggregation of the p53 protein, resulting in the loss of its tumor suppressor functions and the acquisition of tumorigenic properties, effectively transforming p53 into an oncogene^46–48^. Despite this understanding, the precise mechanisms driving the development of HGSOC remain unclear. HGSOC originates from the fallopian tube epithelium, and current evidence suggests that its progression follows a stepwise pattern, beginning with early p53 signature lesions that evolve into STICs, which eventually transform into HGSOC^26,27,36^. We found that p53 is upregulated in all types of lesions collected (**Figure 4A**), from p53 signature to the late-stage HGSOC, while there is a complete absence of expression in the surrounding normal tissue (**Figure 2 and Supplementary Figures 1-7**). Instead, misfolded/aggregated p53 is present only in advanced stages of the progression such as STICs and HGOSCs (**Figure 4, Figure 2 and Supplementary Figures 1-7**). This is in line with previous reports detailing p53 aggregation in HG-SOC using different methods^9,16,21,49^.

p53 missense mutations in HGSOC frequently occur in the DNA binding domain (DBD), a region known for its low thermodynamic and kinetic stability^40,50^. Mutations in this domain can exacerbate its inherent instability, leading to protein misfolding, subsequent aggregation, and loss of tumor suppressor functions^9,41^. Sequencing data from three representative patients revealed different missense mutations in the DBD (G226V, C227F, and G245V), confirming the presence of p53 mutations in pre-neoplastic lesions. Notably, the same p53 mutation persisted across different tumor stages, though the extent of p53 aggregation varied, as observed for patient CA9 (**Figure 1**).

In all samples analyzed in this study, we observed a general upregulation of p53 in the nuclei. It has been previously reported that p53 aggregation can occur both in the nucleus and cytosol of HGSOC cells^9,16^. Presence of both cytosolic and nuclear p53 aggregates correlated with worse prognosis and higher resistance to chemotherapy in a small HGSOC patients series^16^. While we did not observe cytosolic p53 expression or aggregation in the present studies, this could be due to the presence of different types of mutations that can lead to different types of p53 accumulation. For instance, none of the cases presented carried either R273H, R248Q and R248W p53 mutants that have all been shown to cause accumulation both in the nucleus and cytosol^16^.

The mechanisms leading to the change of p53 conformation between pre-malignant and malignant cancer are not well understood. Mutated p53 can be regulated by the same signals that regulate wt p53. It has been shown that the absence of MDM2 or p16INK4a in mice leads to the stabilization of mutant p53, resulting in earlier tumor formation and reduced survival compared to mice with only mutant p53^51^. Additionally, certain pathways may specifically affect subgroups of p53 mutants. For example, the FBXO42-CCDC6-USP28 pathway acts as a positive regulator, enhancing the stability of mutants p53, while C16orf72/HAPSTR1 act as a negative regulator that affects the stability of both wt and mutant p53, depending on its localization and the activity of the E3 ubiquitin ligase HUWE1^52^. The accumulation of p53 can also be facilitated by the interaction with chaperones, like Hsp70 and Hsp90^53,54^. Once p53 aggregates, it can sequester other forms of p53, both mutated and wildtype, but also other proteins like the cofactors p63 and p73, inactivating their functions and promoting the upregulation of antiapoptotic and proliferative factors^41^ which can favor tumor progression. An investigation of stomach and oral cancer biopsies found a correlation between p53 aggregation and tumor stage^55^. The study showed that the levels of aggregated p53 increased in more advanced tumors, and further support the hypothesis that aggregated p53 can be a contributing factor to tumor progression^55^.

Our data suggest that p53 misfolding and aggregation is a structural defect that can distinguish between pre-malignant and malignant tissue in ovarian cancer and thus may play a role in the development of HGSOC. This insight could provide a new perspective on how early-stage lesions progress to invasive cancer and might offer a potential target for early chemopreventive intervention. This can be particularly relevant for patients with mutations in BRCA1/2, who have an increased risk of developing HGSOC and often present early lesions well before full tumor progression^26,56,57^ yet may choose to delay preventative oophorosalpingectomy procedures until menopause^58,59^. These patients could benefit from preventive therapy that blocks progression to malignancy before undergoing surgery^36,56^. Understanding the role and the mechanisms that regulate p53 aggregation will facilitate the development of new drugs aimed specifically at targeting this conformational defect^17^.

## Acknowledgments

We thank the UCLA Translational Pathology Core Laboratory (TPCL) for assistance with histology and the Department of Medicine Statistics Core (DOMStat) for the help in the statistical analysis. We would like to thank Dr. Danielle K. Manning at CAMD core at Dana-Farber Cancer Institute for assistance in targeted sequencing of tissue and Dr. Sanaz Memarzadeh for help accessing archival de-identified cases. This work was supported by the NIH R01CA240718 grant to AS. We acknowledge additional support from the Marsha Rivkin Foundation (Lynda’s Fund Award to AS) and the Iris Cantor UCLA Women’s Education Research Center (to AS).

## MATERIALS & METHODS

### Patient samples collection

De-identified archival tissue specimens were obtained through the Translational Pathology Core Laboratory (TPCL) at UCLA. Fallopian tubes tissues from ten patients were obtained from salpingo-oophorectomies performed on sporadic High Grade Serous Ovarian Cancer cases or from preventive procedures in BRCA-carriers. Paraffin embedded patient tissues were sectioned and stained with hematoxylin and eosin (H&E). All cases collected were reviewed by a gynecologic pathologist that confirmed the diagnosis of p53 signature, STIL, STIC and HGSOC.

### Immunohistochemistry

Paraffin embedded patient tissue sections were baked at 45°C for 20 min and deparaffinized in xylene followed by washes in ethanol and deionized water. Endogenous peroxidases were blocked with Peroxidazed 1 (Biocare Medical, PX968M) at RT for 5 min. Antigen retrieval was performed in the 2100 Retriver chamber (Electron Microscopy Sciences) using the Diva Decloacker buffer (Bio-care Medical, DV2004LX) at 110°C for 15 min. For p53 staining, sections were blocked with Background Punisher (Biocare Medical, BP974L) at RT for 30 min. Primary antibodies were incubated at 4°C overnight using the following dilutions in DaVinci Green diluent (Biocare Medical, PD900L): anti-p53 (DO-1), 1:200 (Santa Cruz Biotech, sc126) and anti-p53 (Pab240), 1:100 (Invitrogen, 13-4100). Following primary antibody incubation sections were incubated sequentially in MACH 3 Mouse HRP probe (Biocare Medical, MP530L) and MACH 3 Mouse HRP-Polymer (Biocare Medical, MH530L) at RT for 20 min each one. Chromogen development was performed with a Betazoid DAB kit (Biocare Medical, BDB2004L). The reaction was quenched with deionized water. 20% Hematoxylin 1 (Richard-Allan Scientific, 7221) was used for counterstaining. The slides were mounted with Permount media (Fisher Chemical, SP15-100). Images were acquired with the Revolve Upright and Inverted Microscope System (Echo Laboratories).

For Ki-67 staining, after heat-induced epitope retrieval in DIVA buffer, another blocking step with Peroxidazed 1 (Biocare Medical, PX968M) at RT for 2 min was performed. Sections were blocked with Background punisher (Biocare Medical, BP974L) for 5 min at RT. The Ki-67/caspase-3 (Biocare Medical, PPM240DSAA) is a prediluted solution and was added to the sample overnight at 4°C. Secondary antibody staining was performed with MACH 2 Double Stain 2 (Biocare Medical, MRCT525H) at RT for 40 minutes. Chromogen development and Hematoxylin 1 staining were performed as previously described.

For PAX8 staining, after heat-induced epitope retrieval in DIVA buffer, another blocking step with Peroxidazed 1 (Biocare Medical, PX968M) at RT for 4 min was performed. Sections were blocked with Background punisher (Biocare Medical, BP974L) for 30 min at RT. Primary antibody PAX8 (Proteintech, 60145-4-Ig) was diluted 1:300 in DaVinci Green buffer (Biocare Medical, PD900L) and incubated overnight at 4°C. Secondary antibody staining was performed with MACH 3 Mouse HRP Probe (Biocare Medical, MP530L) and MACH 3 Mouse HRP-Polymer (Biocare Medical, MH530L) at RT for 15 min each one. Chromogen development and Hematoxylin 1 staining were performed as previously described.

### Lesion images acquisition and quantification

All the slides were scanned using Aperio ScanScope AT whole slide scanner at 20X magnification (Translation Pathology Core Laboratory at UCLA). QuPath 0.3 software was used for digital image analysis, to find and collect lesions for each sample. Images were collected at 20X magnification at high resolution (5000×3560 pixels). The quantification analysis was performed using Fiji Macro tool^45^ (ImageJ 2.1.0), creating custom macro to automatically run the analysis. A color deconvolution (H-DAB type) was performed on each image collected, to split the original color image into three separate channels images: blue, brown, and green. For our analysis we were interested in blue and brown channels that correspond to the hematoxylin and DAB staining, respectively.

The DAB-brown images were enhanced in the contrast (saturation= 0.35) and the threshold (triangle, 0-120) was applied to create a binary image. Nuclei segmentation (watershed) and nuclei dimensions range (500-infinity) were settled.

The Hematoxylin-blue images were enhanced in the contrast (saturation= 0.35) and the threshold (Default, 0-110) was applied to create a binary image. Nuclei segmentation (watershed) and nuclei dimensions range (300-infinity) were settled. The result is the number of nuclei stained with Hematoxylin and DAB in each image. The total number of nuclei per image was obtained from the sum of the total number of nuclei counted with hematoxylin and the total number of nuclei counted in the DAB-stained slide. The positive index was calculated as the ratio between the total number of IHC nuclei on the total number of hematoxylin nuclei. To evaluate the level of p53 aggregation on the total of p53, the positive index was used to calculate the ratio between the Pab240 and DO-1.

### Targeted Sequencing

We selected 10 slides from FFPE tissue blocks for each patient. Samples were sent to the Center for Advanced Molecular Diagnostics (CAMD) at Brigham and Women’s Hospital for sequencing and analysis selected areas using the OncoPanel v3. The results are plotted as Circos plots using R package RCircos^60^.

## Supplementary Material

## Supplementary Table

**Supplementary Table 1.** Lesions collected and tumor progression for each patient. Summary of lesions collected for each patient, used to monitor the progression of the tumor. We aimed to collect about 10 examples of each lesion in each patient whenever possible. + indicates that the collected lesions are between 1 and 5, ++ indicates that the collected lesions are between 5 and 10. p53sign: p53 signature.

| Patient ID | p53sign | STIL | STIC | HGSOC |
| --- | --- | --- | --- | --- |
| CA1 | + | + | + |  |
| CA2 |  |  | ++ | ++ |
| CA3 |  |  |  |  |
| CA4 | ++ | ++ | ++ | ++ |
| CA5 | + | ++ | + | + |
| CA6 | + | ++ | ++ | ++ |
| CA7 | + | + | + | + |
| CA8 | + | ++ | + | + |
| CA9 | ++ | ++ | ++ | ++ |
| CA10 | + | + | + |  |

**Supplementary Table 2.** Adjusted p-value for each comparison Pab240 / DO-1. P-values obtained for each patient by performing a Kruskal-Wallis test with Dunn’s correction for multiple comparisons. NA, data not available. The p-values for the Early lesions vs Late lesions comparison were obtained by performing a Mann-Whitney two-tailed pairwise comparison. NA, data not available. p53sign: p53 signature.

|  | p53sign vs. STIL | p53sign vs. STIC | p53sign vs. HGSOC | STIL vs. STIC | STIL vs. HGSOC | STIC vs. HGSOC | Early vs Late |
| --- | --- | --- | --- | --- | --- | --- | --- |
| CA1 | >0.9999 | 0.4406 | NA | >0.9999 | NA | NA | 0.2619 |
| CA2 | NA | NA | NA | NA | NA | >0.9999 | NA |
| CA4 | >0.9999 | 0.061 | <0.0001 | 0.978 | 0.0065 | 0.5234 | <0.0001 |
| CA5 | >0.9999 | 0.7165 | 0.1864 | >0.9999 | 0.4501 | >0.9999 | 0.035 |
| CA6 | >0.9999 | 0.7442 | 0.6763 | >0.9999 | >0.9999 | >0.9999 | 0.1007 |
| CA7 | >0.9999 | >0.9999 | 0.5514 | >0.9999 | 0.916 | >0.9999 | 0.0952 |
| CA8 | 0.916 | 0.3617 | 0.6054 | 0.0619 | 0.0898 | >0.9999 | 0.0007 |
| CA9 | >0.9999 | 0.0014 | 0.0345 | 0.0011 | 0.0202 | >0.9999 | <0.0001 |
| CA10 | 0.571 | 0.0674 | NA | >0.9999 | NA | NA | 0.0893 |

## Supplementary Figures

**Supplementary Figure 1.**
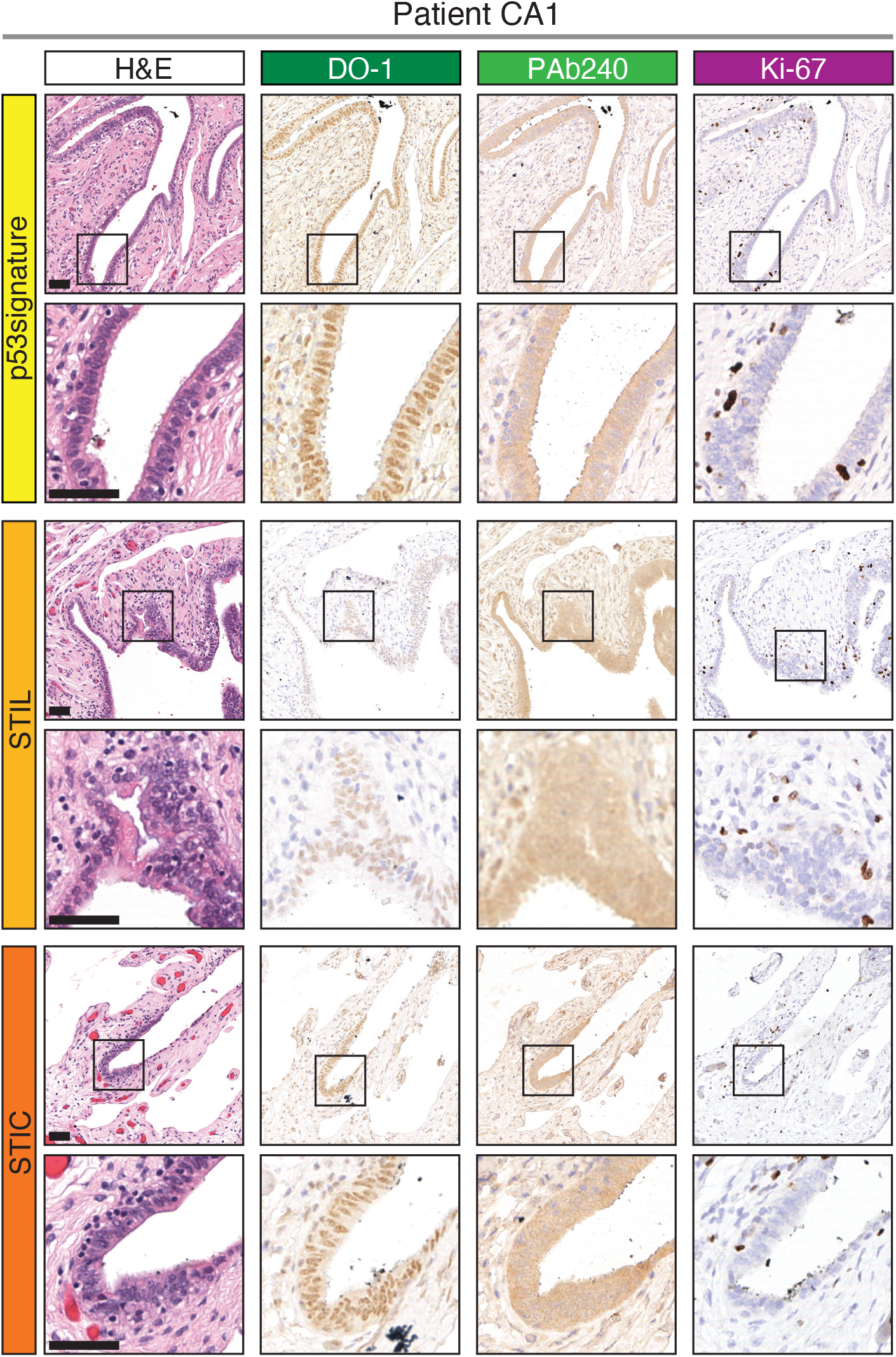
Representative lesions areas for patient CA1. Staining with H&E, DO-1, PAb240 and Ki-67 are reported for each representative lesion. Early lesions are characterized by p53 upregulation by not p53 aggregation. Black boxed area is magnified on the bottom. Scale bars, 50 µm. magnification 20 µm.

**Supplementary Figure 2.**
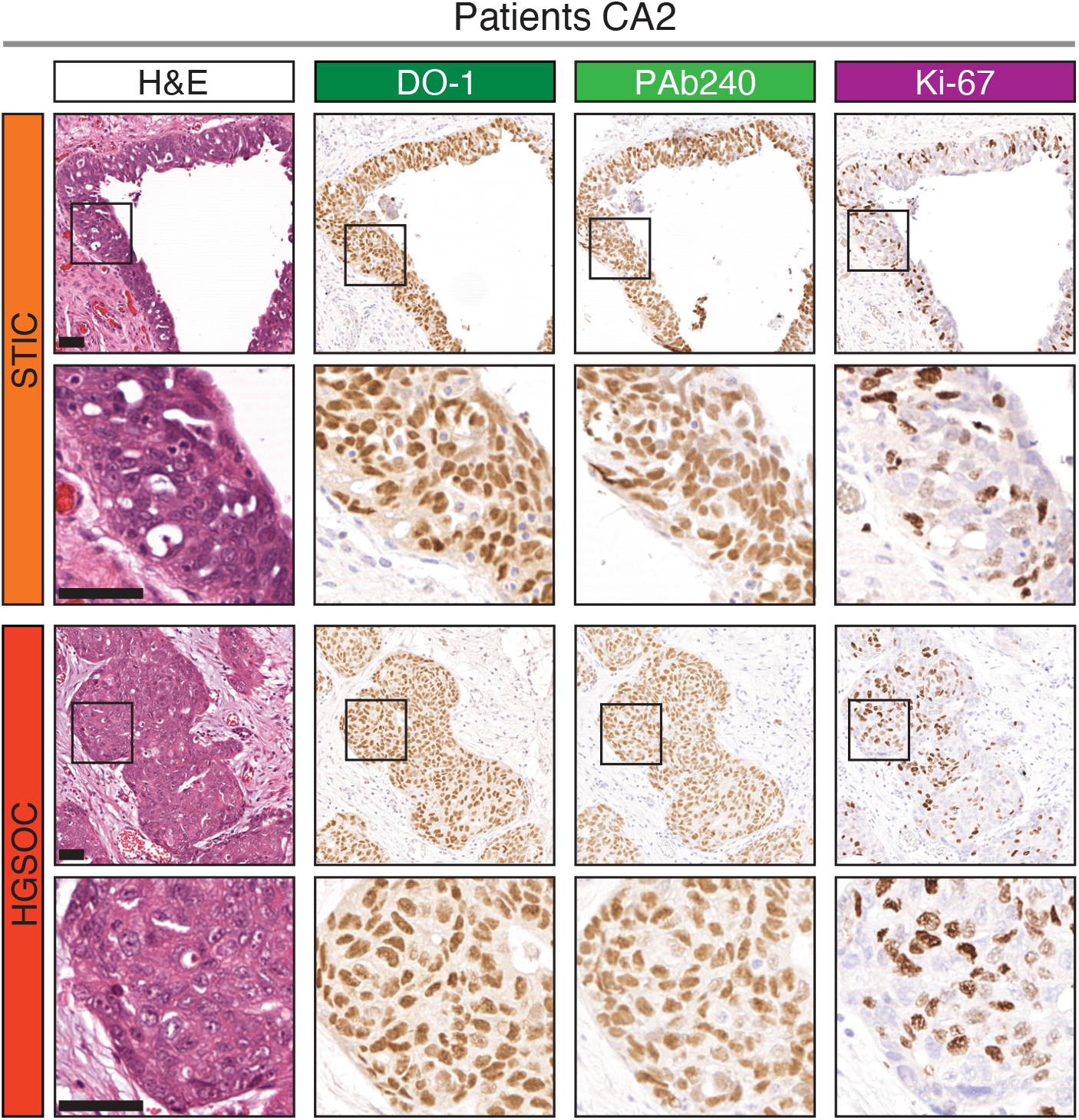
Representative lesions areas for patient CA2. Staining with H&E, DO-1, PAb240 and Ki-67 are reported for each representative lesion. Advanced lesions are characterized by positive PAb240 staining. Black boxed area is magnified on the bottom. Scale bars, 50 µm. magnification 20 µm.

**Supplementary Figure 3.**
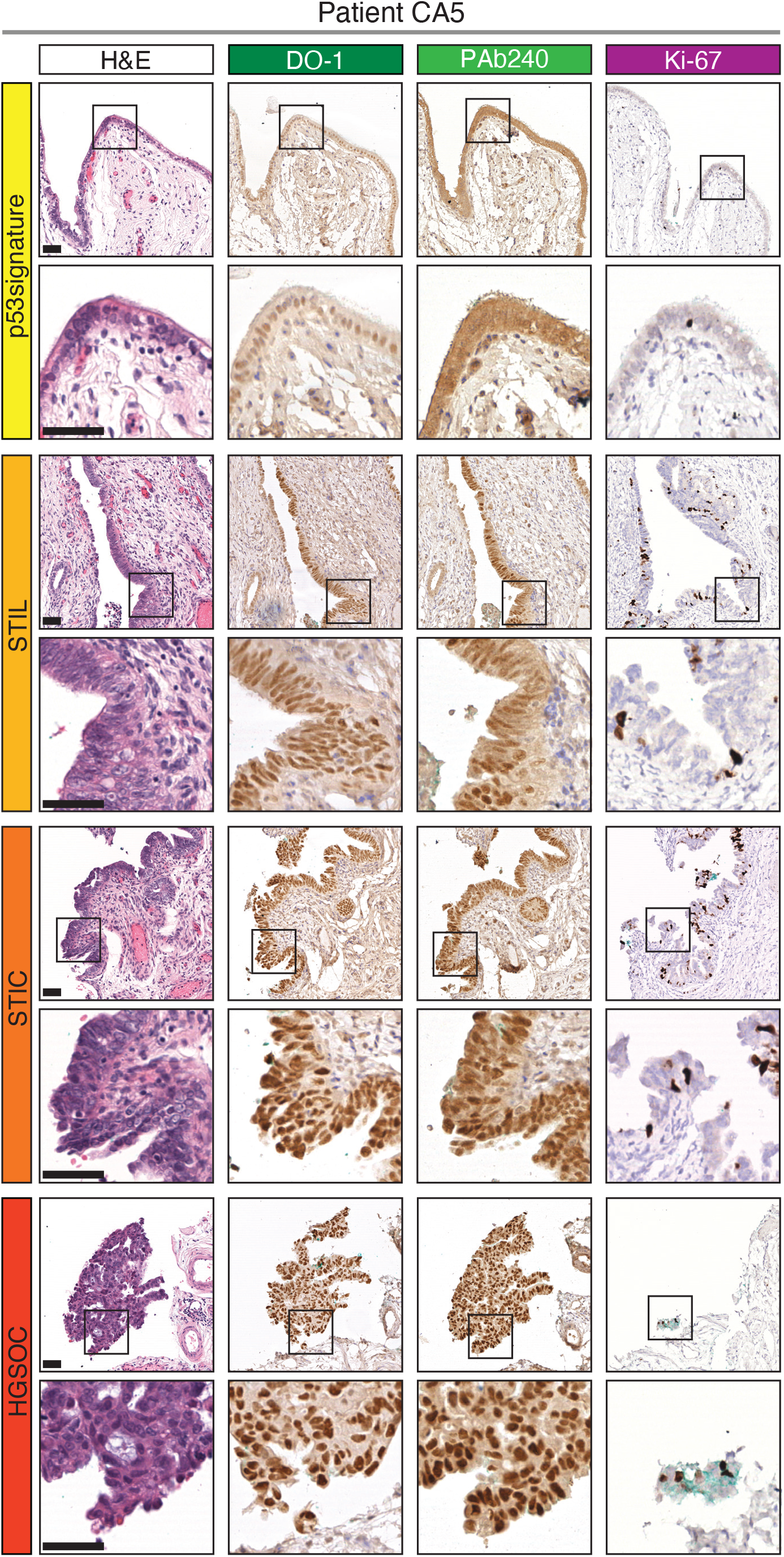
Representative lesions areas for patient CA5. Staining with H&E, DO-1, PAb240 and Ki-67 are reported for each representative lesion. Advanced lesions are characterized by positive PAb240 staining, while early lesions show only non-specific background. Black boxed area is magnified on the bottom. Scale bars, 50 µm. magnification 20 µm.

**Supplementary Figure 4.**
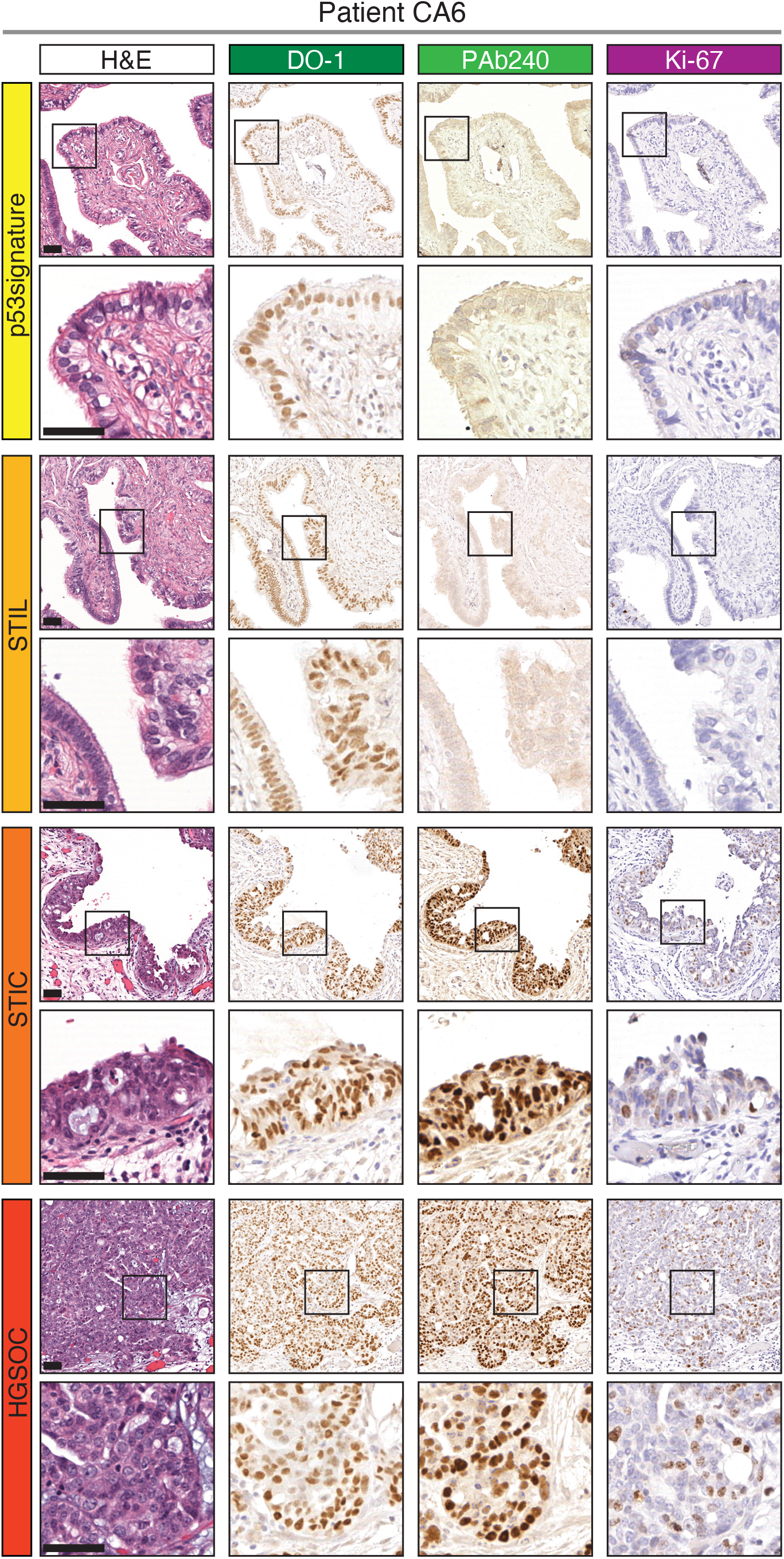
Representative lesions areas for patient CA6. Staining with H&E, DO-1, PAb240 and Ki-67 are reported for each representative lesion. Advanced lesions are characterized by positive PAb240 staining, while early lesions show only non-specific background. Black boxed area is magnified on the bottom. Scale bars, 50 µm. magnification 20 µm.

**Supplementary Figure 5.**
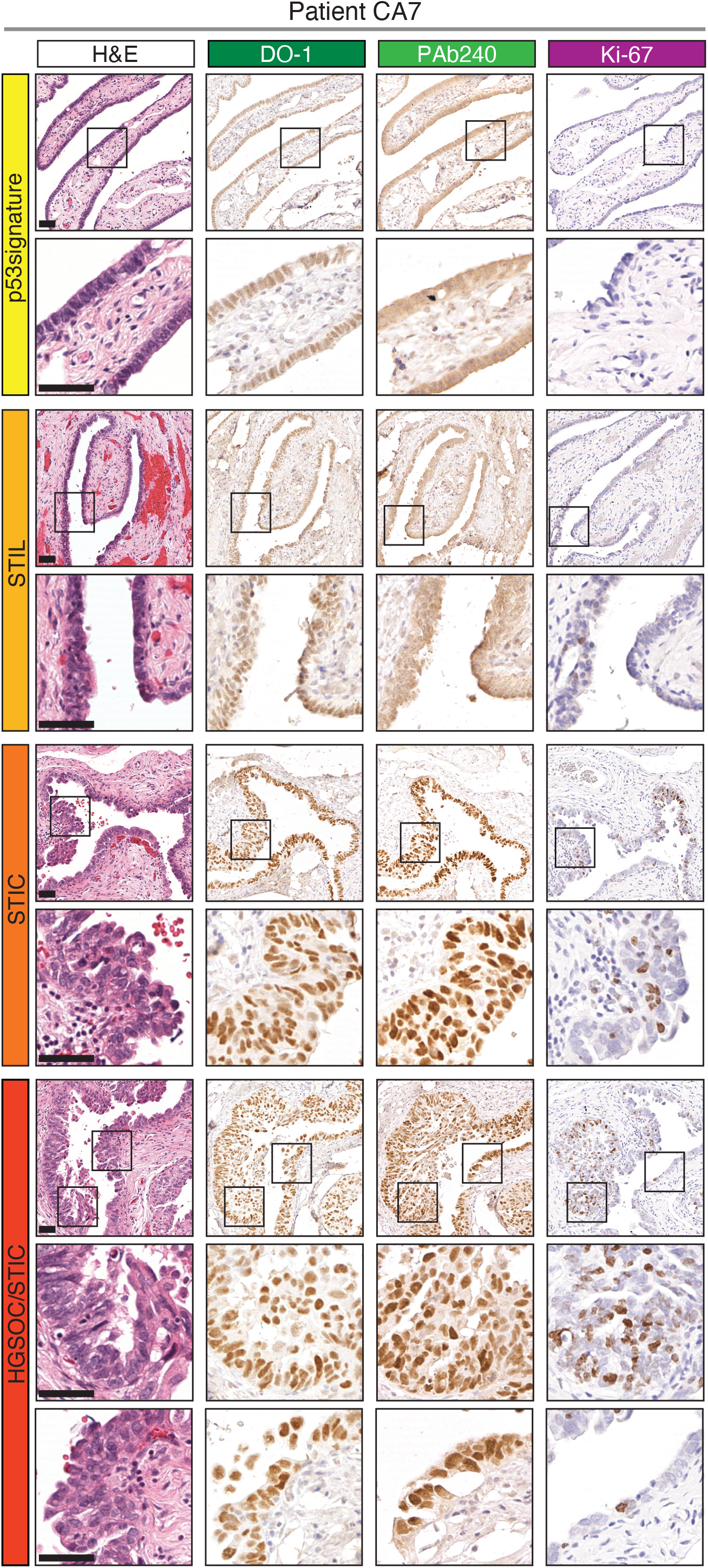
Representative lesions areas for patient CA7. Staining with H&E, DO-1, PAb240 and Ki-67 are reported for each representative lesion. Advanced lesions are characterized by positive PAb240 staining, while early lesions show only non-specific background. Black boxed area is magnified on the bottom. Scale bars, 50 µm. magnification 20 µm.

**Supplementary Figure 6.**
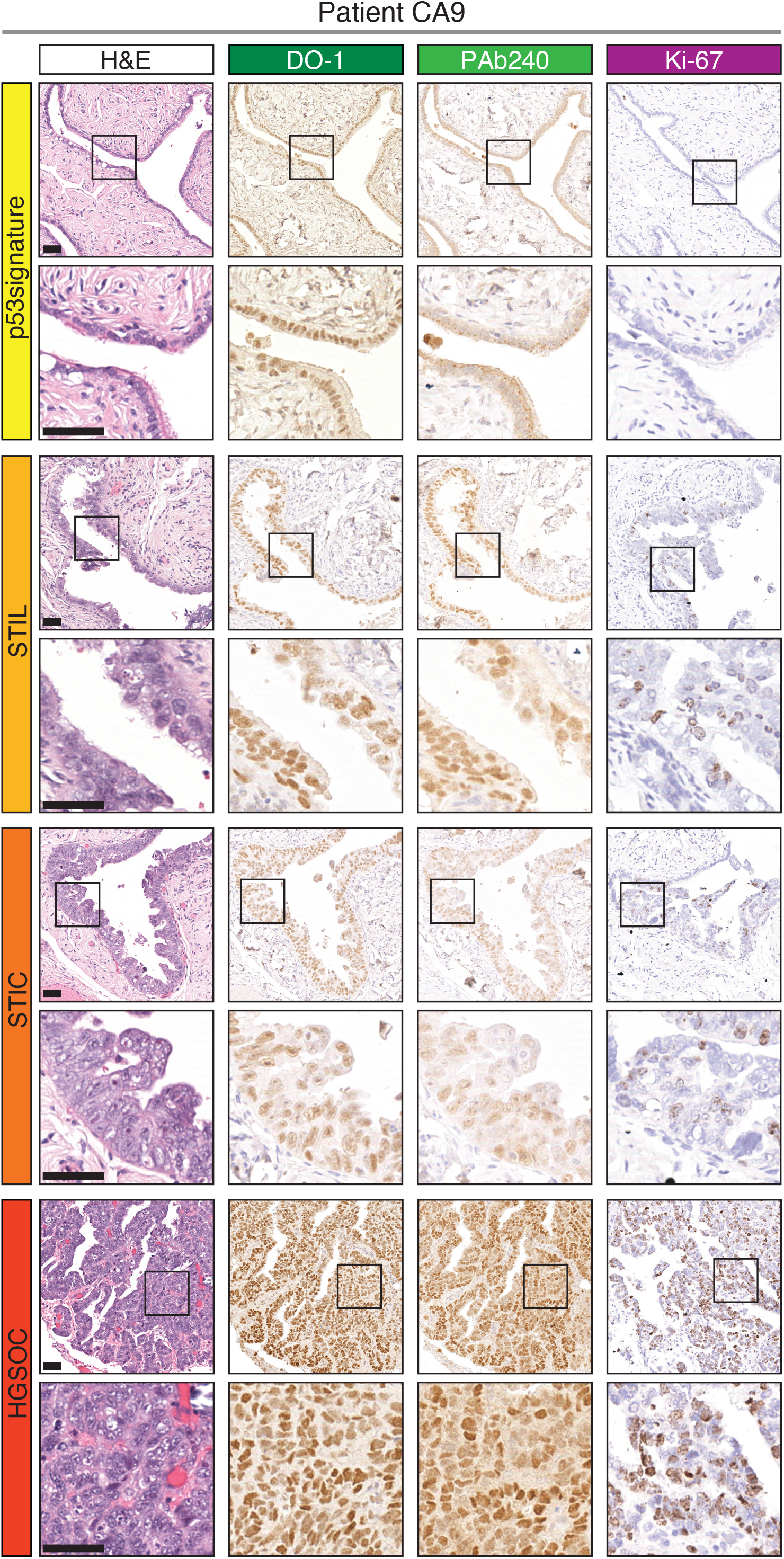
Representative lesions areas for patient CA9. Staining with H&E, DO-1, PAb240 and Ki-67 are reported for each representative lesion. Advanced lesions are characterized by positive PAb240 staining, while early lesions show only non-specific background. Black boxed area is magnified on the bottom. Scale bars, 50 µm. magnification 20 µm.

**Supplementary Figure 7.**
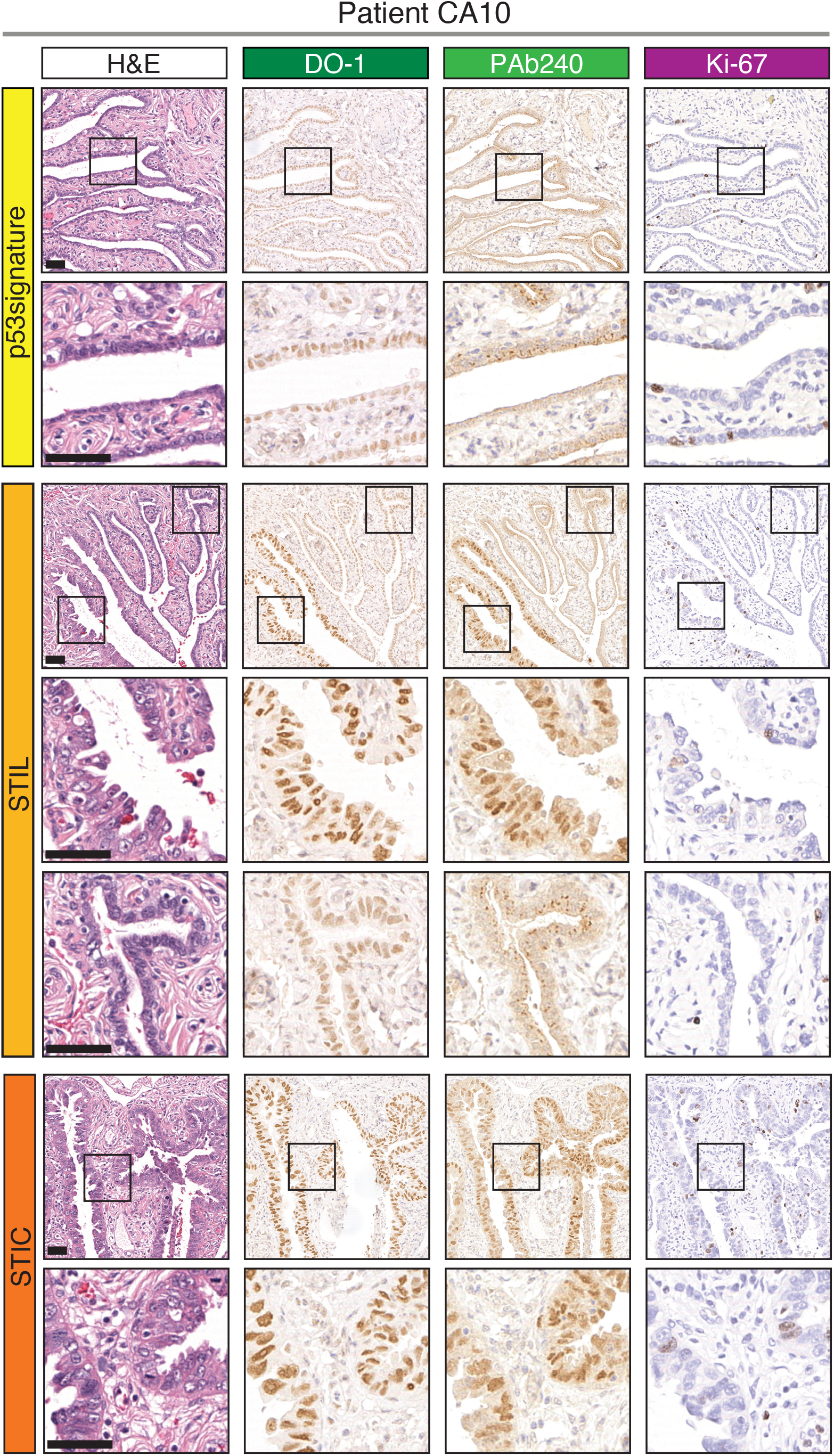
Representative lesions areas for patient CA10. Staining with H&E, DO-1, PAb240 and Ki-67 are reported for each representative lesion. Advanced lesions are characterized by positive PAb240 staining, while early lesions show only non-specific background. Black boxed area is magnified on the bottom. Scale bars, 50 µm. magnification 20 µm.

**Supplementary Figure 8.**
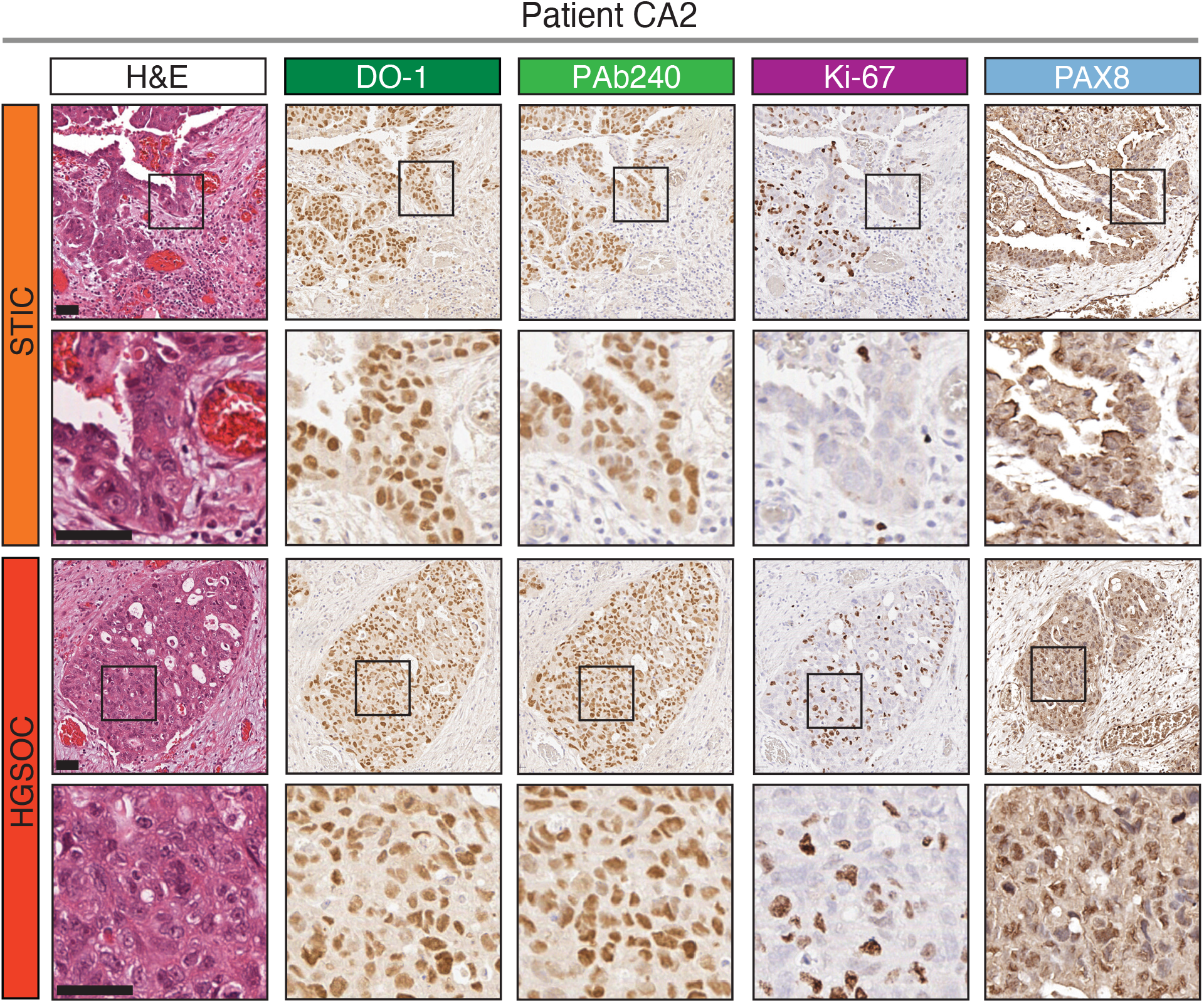
Pax8 staining indicate upregulation in late stages. Representative areas for patient CA2. Staining with H&E, DO-1, PAb240, Ki-67 and Pax8 are reported for each representative lesion. Pax8 is strongly expressed in STIC and HGSOC. The upregulation of Pax8 is found in the same regions where p53 is upregulated. The black boxed area is magnified on the bottom. Scale bars, 50 µm, magnification 20 µm.

**Supplementary Figure 9.**
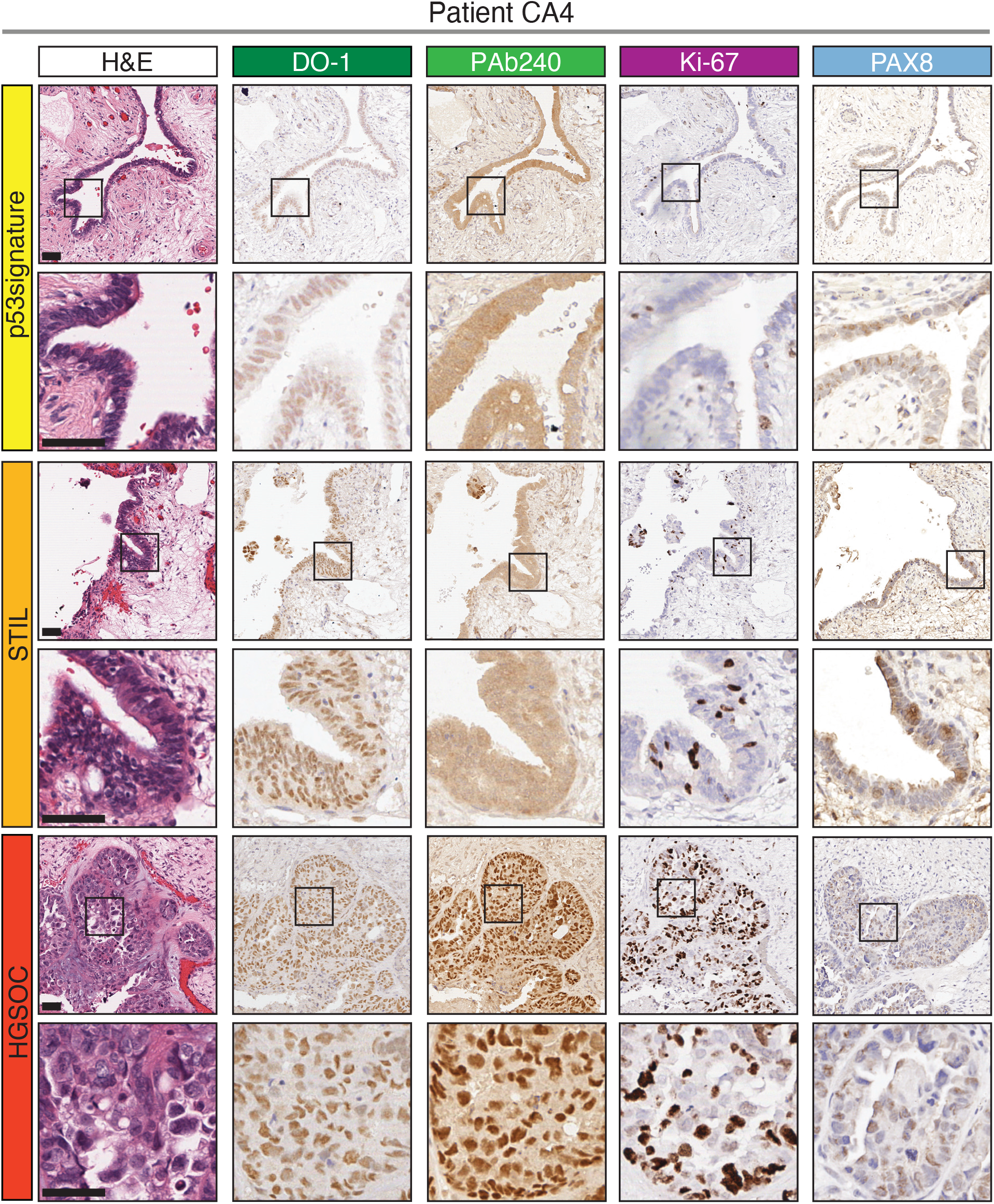
Pax8 staining indicate upregulation in late stages. Representative areas for patient CA4. Staining with H&E, DO-1, PAb240, Ki-67 and Pax8 are reported for each representative lesion. Pax8 is strongly expressed in STIC and HGSOC, while in p53signature and STIL only some nuclei are positive. The upregulation of Pax8 is found in the same regions where p53 is upregulated. The black boxed area is magnified on the bottom. Scale bars, 50 µm, magnification 20 µm.

**Supplementary Figure 10.**
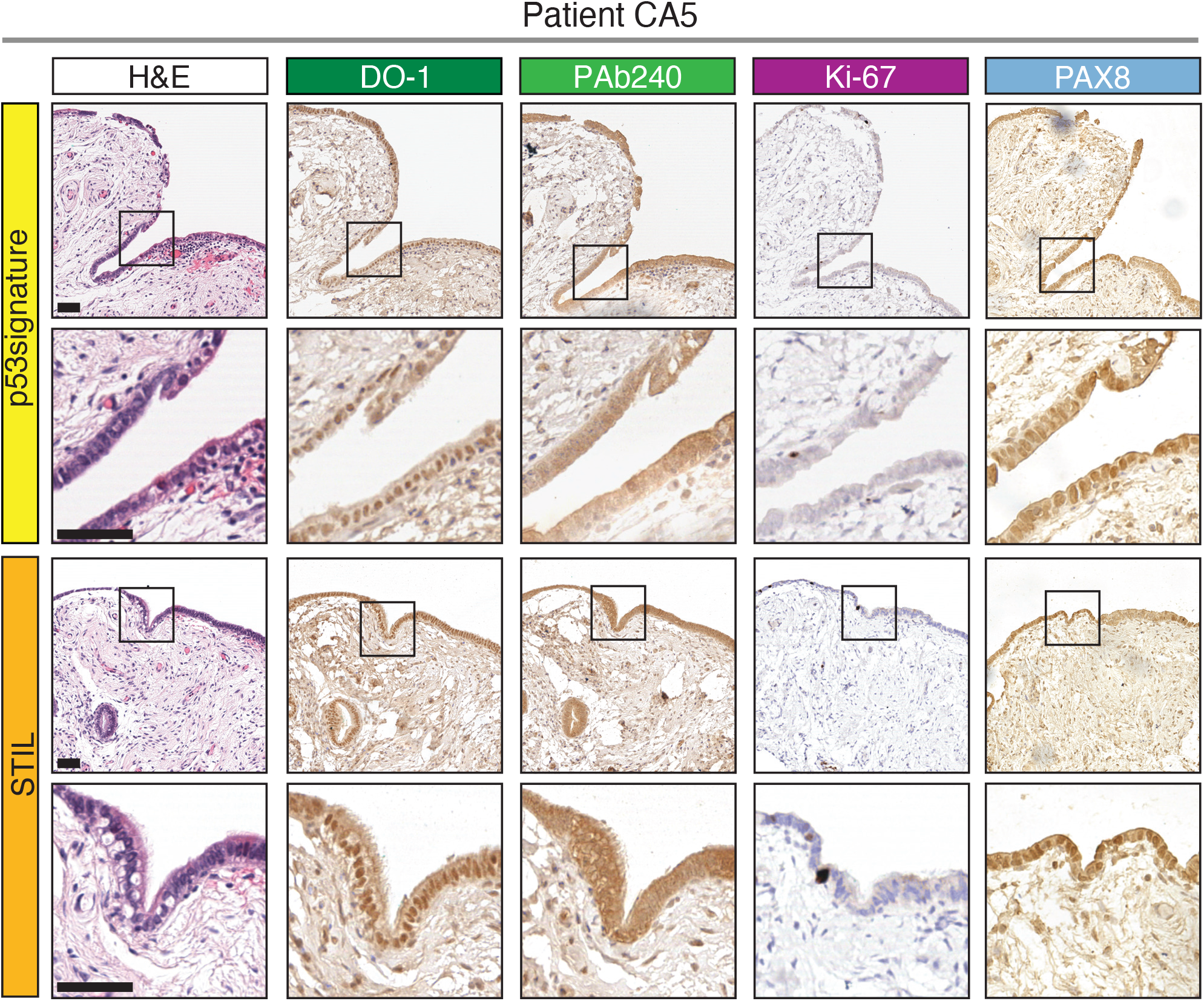
Pax8 staining indicate upregulation in late stages. Representative areas for patient CA5. Staining with H&E, DO-1, PAb240, Ki-67 and Pax8 are reported for each representative lesion. Pax8 staining in p53signature and STIL show only some positive nuclei. The upregulation of Pax8 is found in the same regions where p53 is upregulated. The black boxed area is magnified on the bottom. Scale bars, 50 µm, magnification 20 µm.

**Supplementary Figure 11.**
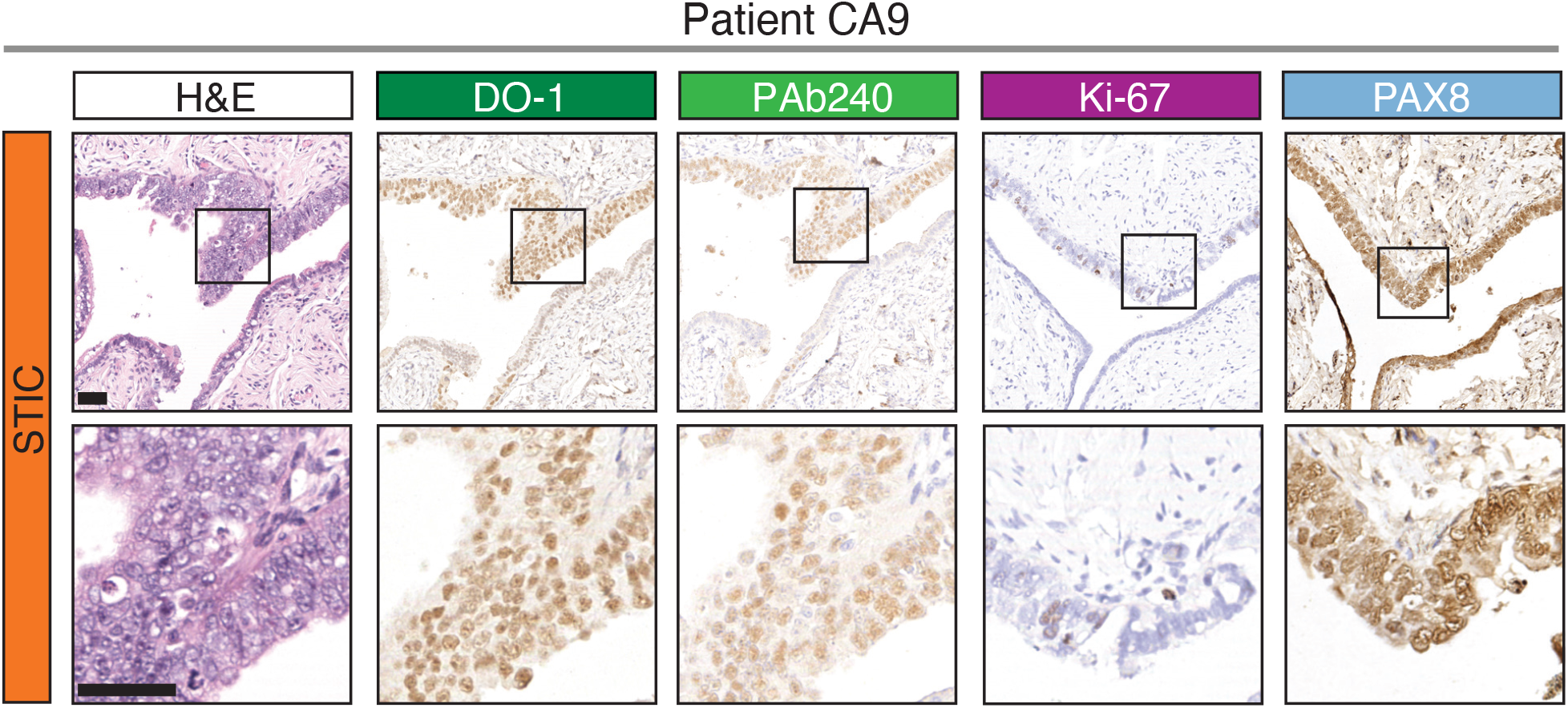
Pax8 staining indicate upregulation in late stages. Representative areas for patient CA9. Staining with H&E, DO-1, PAb240, Ki-67 and Pax8 are reported for each representative lesion. Pax8 is strongly expressed in STIC. The upregulation of Pax8 is found in the same regions where p53 is upregulated. The black boxed area is magnified on the bottom. Scale bars, 50 µm, magnification 20 µm.

**Supplementary Figure 12.**
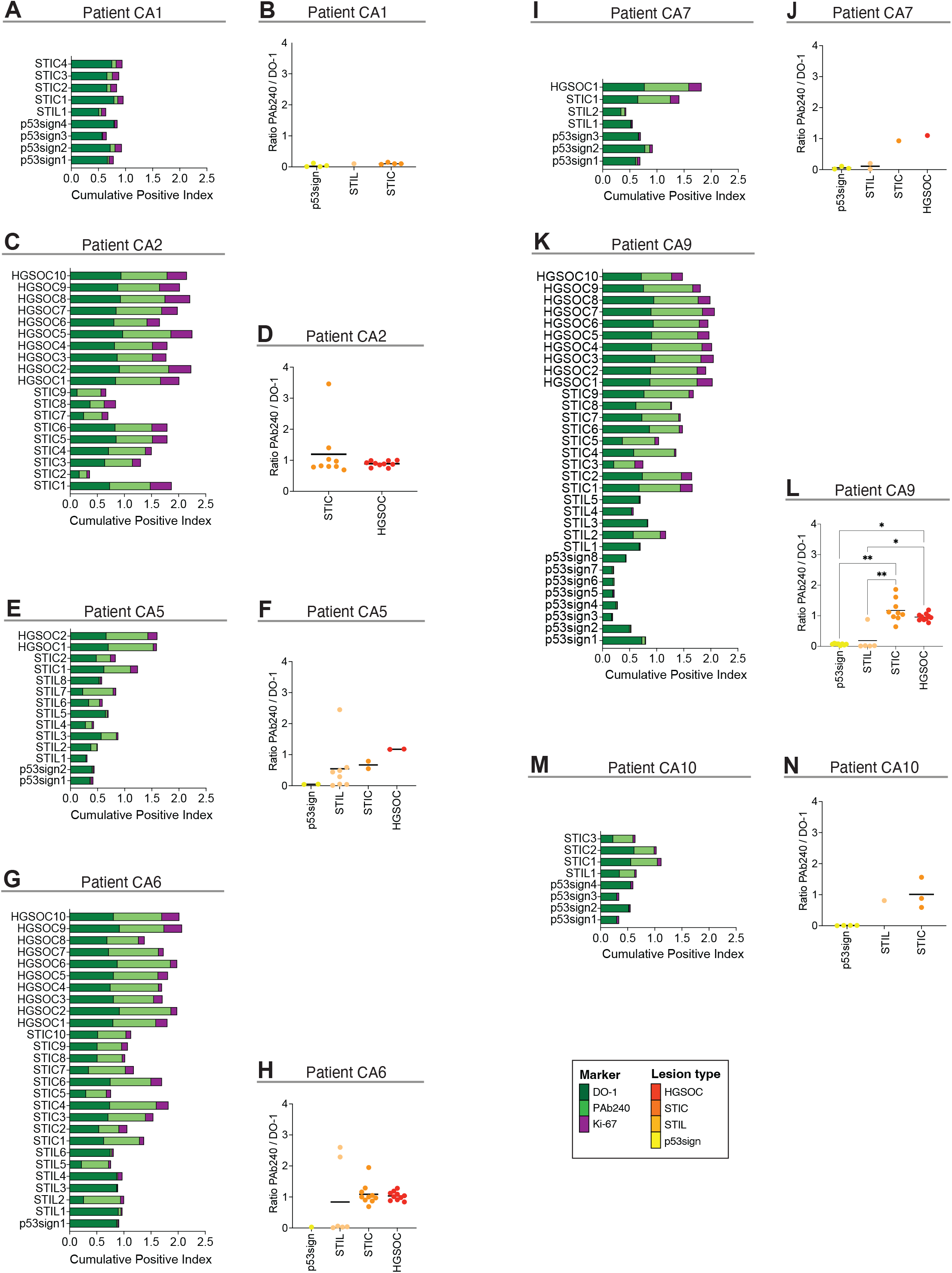
Quantification of total and aggregated p53 in all patients. (A-C-E-G-I-K-M) Positive index calculated for DO-1 (light green), PAb240 (dark green) and Ki-67 (purple) is reported for each lesion collected for all the patients. Each bar represents a lesion. The graphs report the cumulative positive index and show that aggregation of p53 is present only in late stages of the tumor progression. p53sign: p53 signature. (B-D-F-H-L-N) Ratio of Pab240 and DO-1 calculated for each lesion (p53 signature yellow, STILs light orange, STIC dark orange, HGSOC red). CA1 (B): p53sign n= 4, STIL = 1, STIC n= 4; CA2 (D): STIC n= 9, HGOSC n=10; CA5 (F): p53sign n=2, STIL n= 8, STIC n= 2, HGSOC n= 2; CA6 (H): p53sign n= 1, STIL n= 6, STIC n= 10, HGSOC n= 10; CA7 (J): p53sign n= 3, STIL n= 2, STIC n= 1, HGSOC n=1; CA9 (L): p53sign n= 8, STIL n= 5, STIC n= 9, HGOSC n= 10; CA10 (N): p53sign n= 4, STIL n= 1, STIC n=3. Statistical significance tested by performing a Kruskal-Wallis test with Dunn’s correction for multiple comparisons. *p < 0.05, **p < 0.05; Statistical analysis for CA2 was done by performing a Mann-Whitney two-tailed pairwise comparison. ns are not specified in the graphs.

**Supplementary Figure 13.**
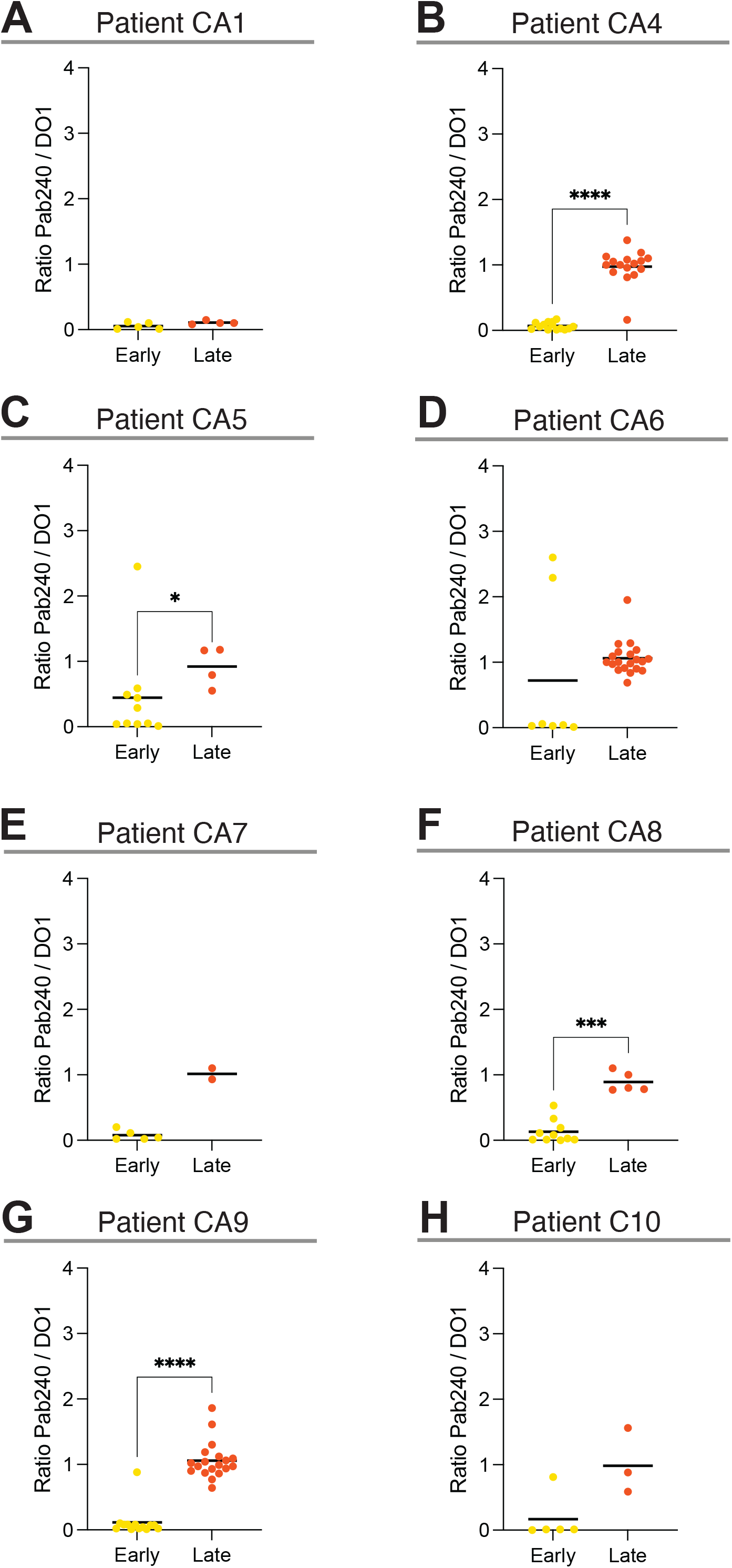
Ratio of Pab240 and DO-1 between early and late lesions. Ratio of Pab240 and DO-1 calculated by pooling together all early lesions (p53 signature and STILs) in yellow, and all late lesions together (STIC and HGSOC) in orange. Each dot represents the ratio of a single lesion. Statistical significance tested by performing a Mann-Whitney test for a two-tailed pairwise comparison. *p < 0.05, ***p < 0.005, ****p < 0.0001; ns are not specified in the graphs. CA1 (A): early n= 5, late n= 4; CA4 (B): early n= 14, late n= 16; CA5 (C): n= 10, late n= 4; CA6 (D): early n= 7, late n= 20; CA7 (E): early n= 5, late n= 2; CA8 (F): early n= 10, late n= 5; CA9 (G): early n= 13, late n= 19; CA10 (H): early n= 5, late n= 3. Patient CA2 is not included in this analysis since it is characterized only by late lesions.

**Supplementary Figure 14.**
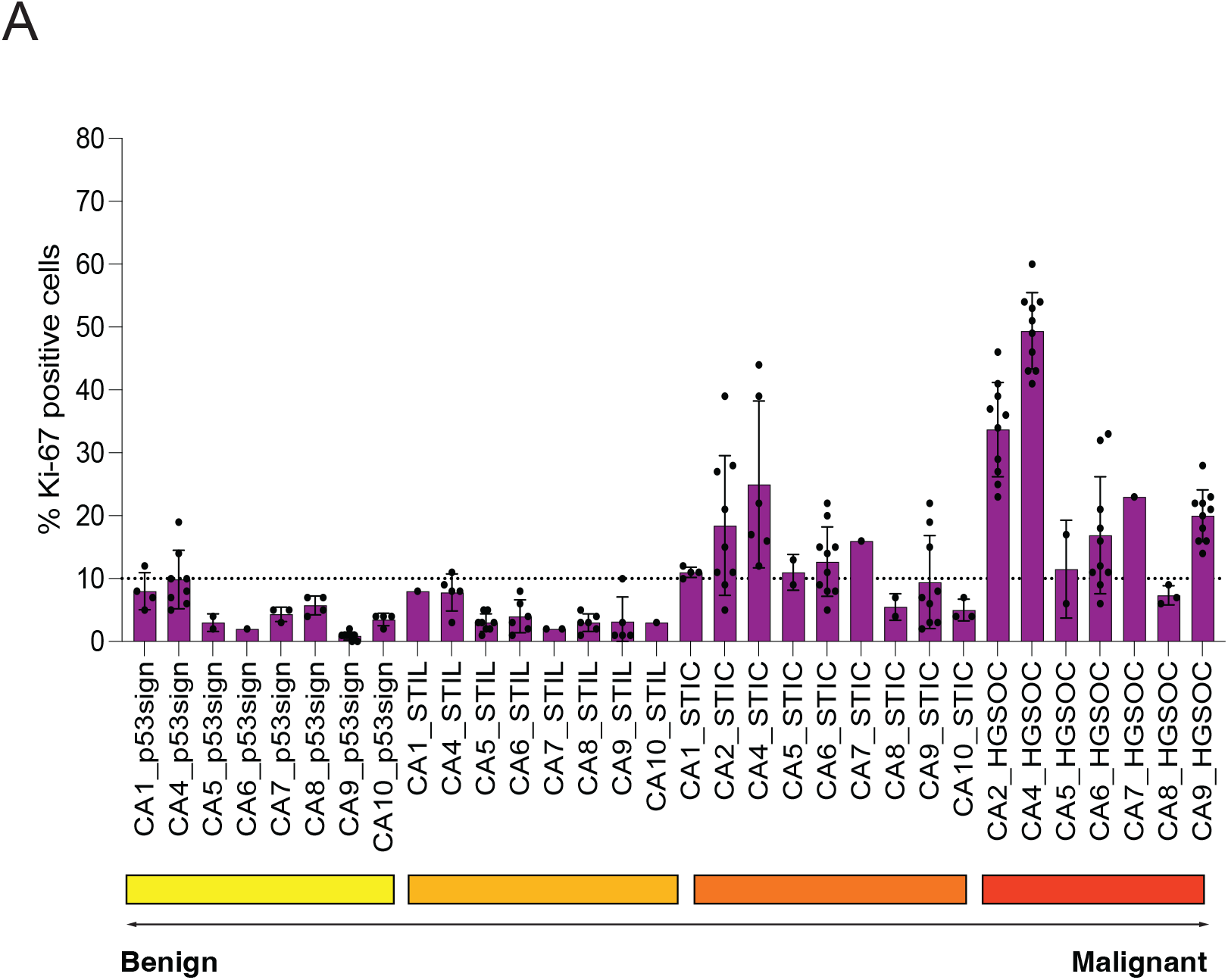
Proliferative index of each patient lesions. Quantification of the proliferative index of each type of lesion in each patient. The graph shows how the proliferative index increases with the progression of lesions from benign to malignant (p53 signature yellow, STILs light orange, STIC dark orange, HGSOC red). The threshold at 10% determines the separation between low proliferative index, distinctive of early lesions and high proliferative index, characteristic of late lesions. Data are organized by lesion progression from benign to malignant. p53sign: p53 signature.

## Notes

### Competing Interest Statement

The authors have declared no competing interest.

